# Protein Docking and Steered Molecular Dynamics Reveal Alternative Regulatory Sites on the SERCA Calcium Transporter

**DOI:** 10.1101/2019.12.19.883355

**Authors:** Rebecca F. Alford, Nikolai Smolin, Howard S. Young, Jeffrey J. Gray, Seth L. Robia

## Abstract

The transport activity of the calcium ATPase SERCA is modulated by an inhibitory interaction with a 52-residue transmembrane peptide, phospholamban (PLB). Biochemical and structural studies have revealed the primary inhibitory site on SERCA, but PLB has been hypothesized to interact with alternative sites on SERCA that are distinct from the inhibitory site. The present study was undertaken to test these hypotheses and explore structural determinants of SERCA regulation by PLB. Steered molecular dynamics (SMD) and membrane protein-protein docking experiments were performed to investigate the apparent affinity of PLB interactions with candidate sites on SERCA. We modeled the relative binding of PLB to several different conformations of SERCA, representing different enzymatic states sampled during the calcium transport catalytic cycle. Overall, the SMD and docking experiments suggest that the canonical binding site is preferred, but also provide evidence for alternative sites that are favorable for certain conformational states of SERCA.

## Introduction

The sarco(endo)plasmic reticulum Ca^2+^-ATPase (SERCA) is responsible for sequestering calcium into the sarcoplasmic and endoplasmic reticulum, creating a reservoir for intracellular Ca^2+^ signaling. This transporter is important in all cell types, and it is tightly controlled in cells where Ca^2+^ handling must rapidly adjust to changing physiological demands (1, 2). A prime example is the regulation of SERCA transport activity in cardiac muscle cells. Calcium transport is reduced under resting conditions in the unstimulated heart through an inhibitory interaction with a single-span transmembrane peptide, phospholamban (PLB), which decreases the affinity of SERCA for its substrate, Ca^2+^. However, inhibition of SERCA by PLB is relieved at high Ca^2+^ concentrations or by phosphorylation of PLB by protein kinase A (PKA) (3) or calcium-calmodulin dependent kinase II (CaMKII) (4, 5) after stimulation of the heart by adrenaline (6). Thus, PLB regulation of SERCA provides an “adrenaline trigger” to increase Ca^2+^ transport and increase cardiac output during exercise. The importance of this regulatory mechanism is underscored by the association of disordered Ca handling and heart disease (7). This makes SERCA an attractive therapeutic target (8, 9) and motivates investigation of the structural elements that govern SERCA function.

Efforts to elucidate the mechanism by which Ca^2+^ and PLB phosphorylation relieve SERCA inhibition have focused on structural changes of the SERCA-PLB regulatory complex. Some studies suggested that SERCA inhibition is relieved by dissociation of the complex (10–12). However, previous studies by our lab (13–16) and others (17–23) indicate that PLB remains bound after phosphorylation and in high Ca, and relief of inhibition is due to a conformational change of the intact regulatory complex. Ca-dependent conformational changes in the SERCA-PLB complex are likely to include alterations in the canonical PLB-binding site on SERCA, a cleft composed of transmembrane helices M2, M6, and M9. Different SERCA crystal structures have captured various enzymatic states of SERCA (24) in which this cleft is relatively more open or closed (**Fig. 1a**). Thus, structural transitions between SERCA conformers are expected to alter contacts with PLB, which may account for our previous observation that the affinity of PLB for SERCA is modestly reduced by Ca binding to SERCA (16). Alternatively, the transitions between enzymatic states (**Fig. 1b**) may result in large-scale reorganization of the SERCA-PLB quaternary structure. In particular, we previously hypothesized that the PLB transmembrane domain could become displaced from the canonical PLB-binding cleft and translocate to a non-inhibitory site on SERCA (16). One possible alternative PLB-binding site is suggested from comparison with a closely related P-type ATPase, the sodium/potassium ATPase (NKA). X-ray crystallography (25, 26) and modeling (27) studies indicated that NKA regulatory subunits, the FXYD family of proteins, bind to the outside of helix M9 (**Fig. 1a, green**). Another accessory binding site was suggested by electron crystallography of the SERCA and PLB, which involves an interaction of PLB with transmembrane segment M3 of SERCA (**Fig. 1a, red**) (28–30). Like the canonical site, the putative novel sites undergo significant structural changes during the SERCA transport cycle (**Fig 1b**).

**Figure 1:**
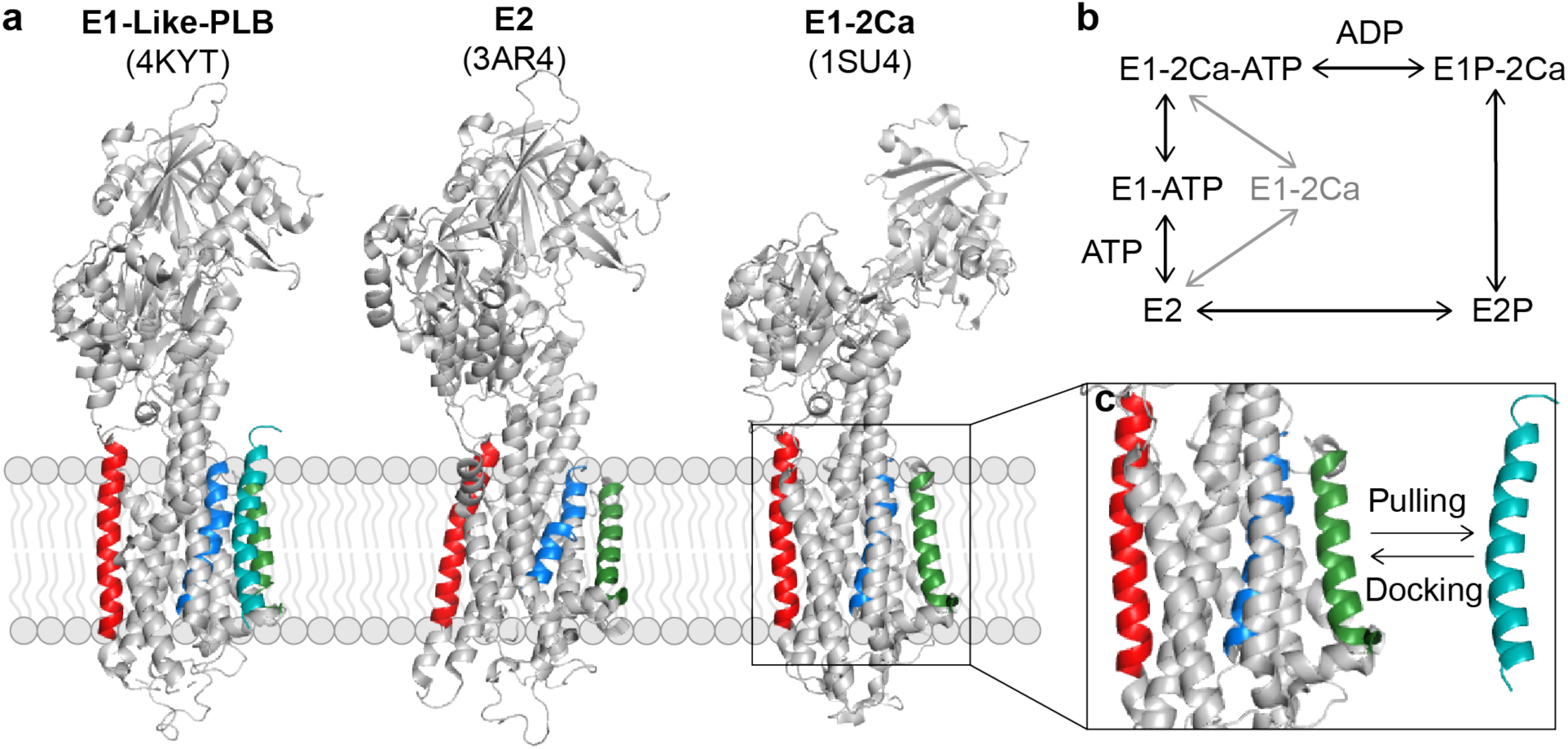
Exploring possible binding sites for phospholamban on different enzymatic states of SERCA. In this study, we examined the interaction of PLB with three different enzymatic states of SERCA. The three states are shown in (a): the E1-like-PLB bound state of SERCA (PDB 4KYT), the E2 calcium-free state (PDB 3AR4), and the E1 calcium bound state (PDB 1SU4). The canonical inhibitory binding site is formed by helices M2, M6 (blue), and M9 (green). The two postulated alternative sites are binding to the outside of helix M9 (green) and M3 (red). The role of each state in the catalytic cycle of ATP-mediated SERCA transport is shown in (b). Panel (c) shows two methods for exploring PLB interactions with SERCA. Steered molecular dynamics simulations pull PLB away from a postulated binding site and quantitate the unbinding force. In contrast, membrane protein docking tests possible interaction sites and quantitates the binding energy.

Here, we used computational tools to determine how PLB might interact with these sites and to identify structural determinants of SERCA regulation. By virtue of their native environment embedded in a lipid bilayer, membrane protein interactions are among the most difficult to measure, model, and simulate. However, the interaction of SERCA with PLB is an excellent membrane protein complex for evaluation by computational structural biology methods. X-ray crystallography has mapped out nearly every step in the SERCA transport cycle (**Fig. 1b**), revealing the different conformations of the intermediate enzymatic states (24, 31). Moreover, a structure that reveals key aspects of the SERCA-PLB complex has been determined by X-ray crystallography (**Fig. 1a, “4KYT”**) (10). This breakthrough was enabled by a mutant form of PLB that greatly increased its affinity and inhibitory potency. The degree to which this mutant recapitulates the regulatory complex structure of the wild-type protein is unknown. In addition, the crystal structure revealed an unexpected E1-like conformation of SERCA bound by PLB. How PLB may interact with other SERCA conformations is another unanswered question.

While the wealth of structural information about the SERCA regulatory complex represents an excellent foundation for analysis, the complexity of the system creates challenges in capturing a full structural description of the regulatory complex. We used several methods to meet the computational challenge of modeling a large complex of more than 1,000 residues embedded in a lipid bilayer environment, while considering numerous conformational states. One approach is steered molecular dynamics (SMD), which can quantify the unbinding force of two proteins in the context of a detailed membrane environment on a timescale accessible to molecular dynamics. SMD captures the membrane environment in atomistic detail; however, it requires the binding location to be designated *a priori*. Further, SMD is computationally intensive, limiting the number of binding sites and modes of binding that can be evaluated. An alternative approach, protein-protein docking, provides a powerful tool to predict the binding location and structure of the bound complex (32, 33). This approach relies on an implicit approximation of the surrounding environment, making it computationally efficient and improving overall scalability. Recently, the development of specialized tools that account for the sampling and chemical properties of the lipid environment has enabled docking of membrane proteins (34). Collectively, these tools provide complementary information for membrane protein-protein docking, enabling us to explore a wide-range of possible binding sites in an unbiased manner while also evaluating feasibility in an atomistic membrane environment. Models from such simulations can provide mechanistic insight to confirm or refute prior hypotheses about the formation of membrane protein complexes.

In the present study, we determined the relative binding of PLB to candidate sites using SMD to quantify the rupture force required to extract PLB docked to a candidate site, with peak force taken as an index of the affinity of PLB for that site (**Fig. 1c, “pulling”**). In addition, we performed an unbiased search for novel candidate sites using membrane protein-protein docking simulations (**Fig. 1c, “docking”**). We modeled the relative binding of PLB to several different conformations of SERCA (**Fig.1a**), representing the major intermediates in the calcium transport cycle (**Fig. 1b**). Together, these techniques provide insight into the structural determinants of the regulatory interaction of PLB with different enzymatic states of SERCA.

## Results

To test whether PLB may bind to putative alternative sites, we performed steered molecular dynamics and protein-protein docking of PLB to different conformations of SERCA. These conformations are sampled as SERCA progresses through the catalytic cycle (**Fig 1B**), binding Ca ions in the cytoplasm and transporting them into the lumen of the sarcoplasmic reticulum. In this study, we examined three key SERCA conformations (**Fig. 1A**): (1) the calcium-free E1-like state bound by PLB (represented by 4KYT) (10), the calcium-free E2 state (represented by 3AR4) (35), which is characterized by a more open PLB-binding groove compared to 4KYT, and (3) the calcium-bound E1 state (represented by 1SU4) (36). The starting structure of the PLB TM domain was taken from the PLB-SERCA complex 4KYT. The cytoplasmic domain of PLB was not detected in this X-ray structure and was omitted from the SMD simulations. Previous biochemical studies have demonstrated that the transmembrane domain is sufficient for binding and inhibition of SERCA (37).

### Steered Molecular Dynamics (SMD) Simulations

To perform SMD simulations, the SERCA residue C*α* atoms positions were first restrained by applying a spring force to each C*α* atom, preventing significant deviations from the starting position. Pulling force was applied uniformly to C-alpha residues of PLB, with force directed away from the center of mass of the SERCA TM domain. **Fig. 2a** quantifies the force applied to PLB (vs. time) for three repeated simulations of PLB interacting with the canonical cleft of 4KYT beginning with the original orientation of PLB in the crystal structure. As the simulation progressed, the number of interacting residues decreased until the interaction was broken. Force increased to a maximum, then rapidly declined to a non-zero plateau. The peak force represents the point at which the PLB-SERCA complex ruptured and was quantified as an index of binding affinity. The plateau force was due to viscous drag of the lipid bilayer after the loss of the protein-protein contacts.

**Figure 2:**
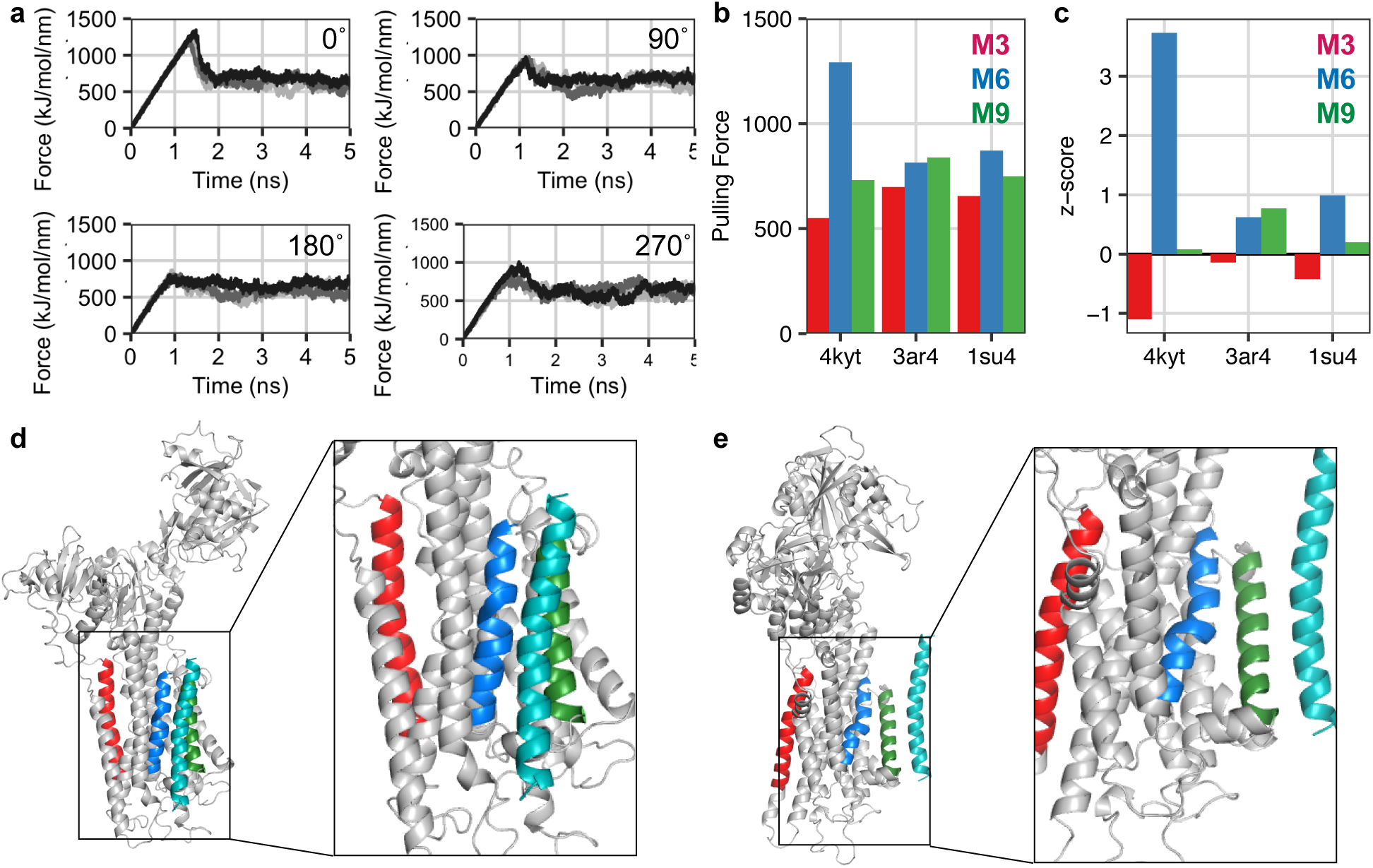
Steered molecular dynamics simulations of PLB interactions with different sites on three enzymatic states of SERCA. (a) Quantification of force that developed as PLB was pulled from the canonical cleft of the E1-like-PLB state of SERCA. Data are three repeated measures of force for the original orientation of the X-Ray crystal structure (PDB 4KYT) and after axial rotation of PLB by 90°, 180°, and 270°. (b) Peak unbinding force at the M3 site (red), the canonical cleft made by M6/M9 (blue), and the site outside M9 (green) on each enzymatic state of SERCA in kJ/mol/nm. (c) Z-score analysis of peak unbinding forces relative to the mean pulling force at the M3 (red), canonical (blue) and outside M9 (green) sites. Structural models of key interactions with non-canonical sites are shown in the bottom row. (d) Interaction of PLB (teal) with the canonical binding site formed by helices M6 (red) and M9 (green) in the E1-calcium bound state of SERCA (PDB 1SU4). (e) Interaction of PLB (teal) with the M9 accessory site (green) in the E2 calcium free state of SERCA (PDB 3AR4).

For the equilibrated PLB-SERCA crystal structure (4KYT), we determined a peak force of 1292 ± 69 kJ/mol/nm (error is standard deviation for n = 3). The simulation was then repeated for PLB rotated around its long axis by 90°, 180°, and 270°. Simulations of alternative axial orientations showed ∼30% lower peak force (**Table 1, Fig. 2b**-**c**), which is consistent with the native orientation of PLB in the X-ray crystal structure being the most stable configuration. Z-score analysis is given in **Supplementary Table S1**.

**Table 1:** Summary of PLB interactions with different SERCA states, quantified by steered molecular dynamics.

| SERCA State | Binding Location | Mean Rupture Force (kJ/mol/nm) |  |  |  |
| --- | --- | --- | --- | --- | --- |
|  |  | 0° | 90° | 180° | 270° |
| E1-like Ca-Free (4KYT) | M3 | 546 ± 53 | 526 ± 41 | 485 ± 38 | 550 ± 27 |
|  | M6/M9 | 1292 ± 69 | 982 ± 3 | 838 ± 31 | 914 ± 84 |
|  | M9 | 686 ± 46 | 731 ± 45 | 688 ± 27 | 616 ± 44 |
| E2 Ca-Free (3AR4) | M3 | 544 ± 20 | 640 ± 52 | 698 ± 14 | 572 ± 9 |
|  | M6/M9 | 848 ± 29 | 814 ± 22 | 729 ± 40 | 720 ± 94 |
|  | M9 | 658 ± 40 | 580 ± 87 | 811 ± 12 | 838 ± 108 |
| E1 Ca-Bound (1SU4) | M3 | 606 ± 49 | 655 ± 23 | 629 ± 25 | 601 ± 39 |
|  | M6/M9 | 871 ± 25 | 775 ± 97 | 833 ± 81 | 766 ± 15 |
|  | M9 | 682 ± 32 | 681 ± 55 | 736 ± 88 | 749 ± 60 |

#### SMD of PLB Docked to Putative Alternative Sites

We performed SMD experiments comparing PLB bound to the canonical M6/M9 site with PLB bound to alternative sites to M3 or to the outside of M9, with three repeated trajectories simulated for each orientation **(Supplemental Fig. S1)**. The average peak force associated with the alternative sites was reduced by approximately 50% compared to the canonical site. Binding to M3 or the outside of M9 was modestly improved by axial rotation of the PLB transmembrane domain by 270° or 90°, respectively (**Table 1; Fig. 2b-c**) but rupture forces were still lower than those observed for various axial orientations of PLB at the canonical site. The data suggest that PLB binds best to the M6/M9 site in the E1-like conformation captured in the SERCA-PLB crystal structure. When bound to the canonical site, PLB interacts with side chains of SERCA helices M4, M6, and M9. In contrast, when docked to the outside of M9 in the manner of binding of FXYD proteins with M9 of the sodium/potassium ATPase (26, 27), PLB had interactions with M9 only which accounts for the lower rupture force of this configuration.

#### SMD of Different SERCA Conformations

Next, we measured the relative binding of PLB to different conformations of SERCA to test how PLB affinity might change as a result of SERCA structural transitions during the catalytic cycle. Repeated SMD simulations revealed that peak rupture force for PLB bound to the canonical binding site (**Fig. 2b-c, Supplementary Fig. S2**) of the calcium-free E2 state (3AR4) structure was 34% lower than that observed for the E1-like state (4KYT). Interestingly, the rupture force for PLB bound to the hypothetical alternative site outside M9 in a 270° orientation (**Fig. 2d**) was slightly improved for E2 compared to the binding of PLB to the outside of M9 of the E1-like structure. Thus, due to improved binding to the outside of M9 and worsened binding to the canonical M6/M9 site, the two alternative sites then showed similar maximal rupture forces (838 ± 108 and 848 ± 29 kJ/mol/nm, respectively). The Ca-bound E1 structure (1SU4) yielded a peak force that was 33% lower than that observed for E1-like state (**Fig. 2e, Supplementary Fig. S3**). E1-Ca also showed a decrease in the rupture force deficit between the M6/M9 site and the proposed site outside M9 compared to the E1-like structure (4KYT). The magnitude of the difference in rupture forces observed for 4KYT (Ca-free) vs. 1SU4 (Ca-bound) is in harmony with our previous physical measurements that suggested a 41% difference in apparent binding for the SERCA-PLB complex in low and high Ca conditions (16). Overall, the data indicate that there is a clear preference for the E1-like PLB-bound state captured by the crystal structure. However, for other enzymatic states, PLB may bind with similar affinity to the canonical site and the putative site on the outside of M9. This result is broadly compatible with the hypothesis of diverse modes of interaction between PLB and SERCA.

### Membrane Protein-Protein Docking of the SERCA-PLB complex

Through steered molecular dynamics simulations, we examined the properties of PLB in four different axial orientations bound to three potential binding sites. To explore the full landscape of potential SERCA-PLB interactions, we pursued a complementary unbiased docking strategy. First, we used fast-Fourier transform (FFT)-based global docking (38) to identify possible binding sites on different enzymatic states of SERCA (represented by structures 4KYT, 3AR4, and 1SU4). For each structure, the FFT method identified 30-50 overall docked SERCA-PLB complexes, ranked by an energy function that uses desolvation and electrostatic energy terms, and filtered based on the requirement for PLB to span the membrane in the correct direction. Using these criteria, we obtained eight SERCA-PLB complexes for the calcium-free E1-like state of SERCA (4KYT) (**Fig. 3a,d**), five for the calcium-free E2 state (3AR4) (**Fig. 3b,e**), and six for the calcium-bound E1 state (1SU4) (**Fig. 3c,f**).

**Figure 3:**
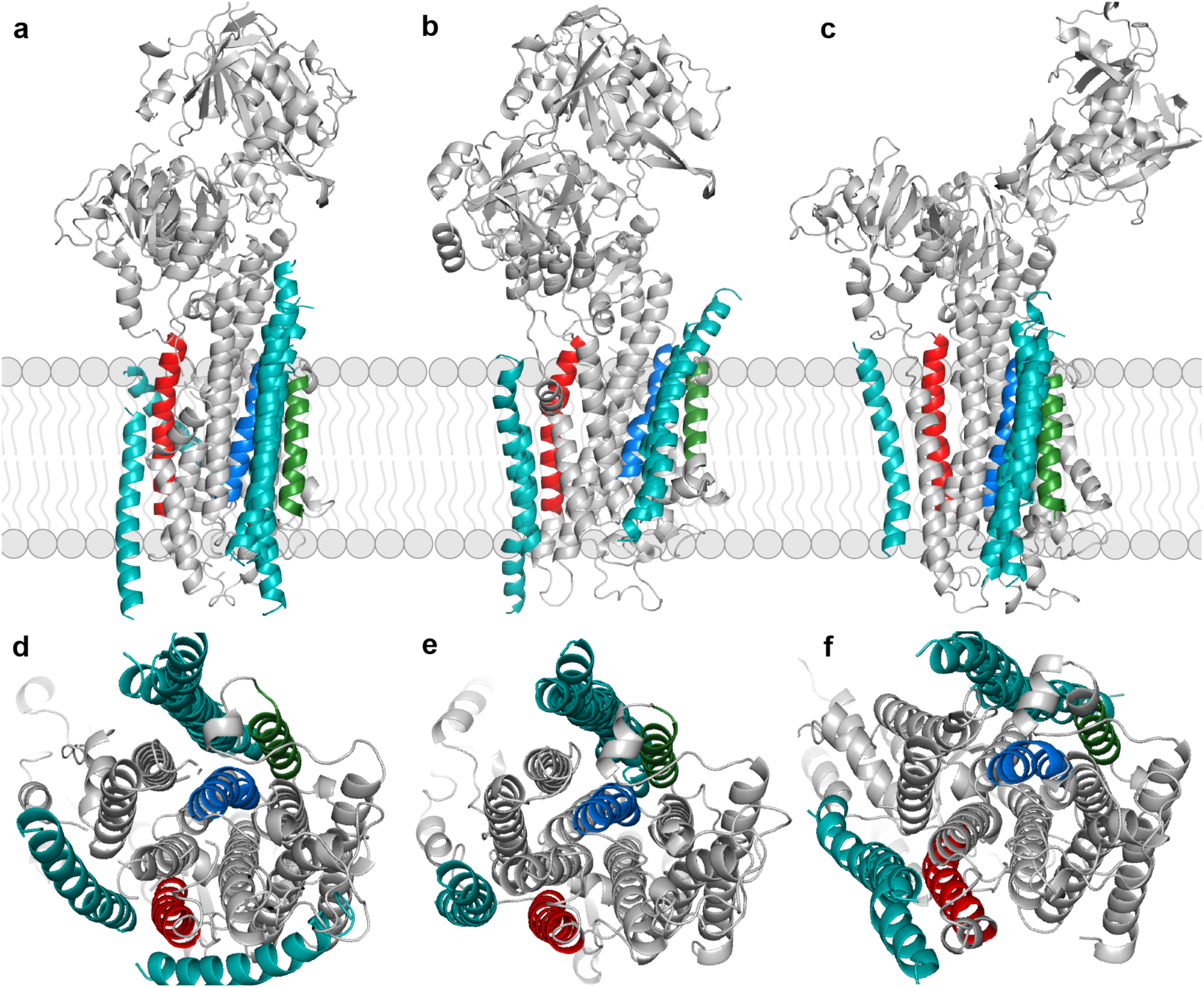
Global protein-protein docking solutions for PLB interaction with different enzymatic states of SERCA. For each enzymatic state of SERCA, we used a fast-Fourier transform (FFT) based global docking program to find possible binding sites and conformations of PLB at those sites. PLB binding solutions (teal) are shown bound to each enzymatic-state of SERCA in a membrane-facing view (top row) and luminal view (bottom row): (a,d) 4KYT, (b,e) 3AR4, and (c,f) 1SU4. The canonical cleft is highlighted with M6 in blue and M9 in green and the alternative sites are highlighted with M3 in red and M9 in green.

Several docking solutions included PLB bound to the M6/M9 canonical site; while the remaining solutions included PLB docked at the M3 accessory site. This result contrasts with the SMD experiments, which showed no appreciable affinity of PLB for this site. Moreover, while the SMD experiments suggested some affinity of PLB for the putative binding site on the outside of helix M9, the unbiased protein-protein docking experiments did not yield M9-docked structures among the most favorable solutions. Additional comparison of the complementary methods is provided in the Discussion section below.

#### Docking of Different Axial Orientations of PLB

After obtaining initial docked SERCA-PLB complexes, we used RosettaMPDock (39) to optimize high-ranking PLB orientations within the membrane environment. RosettaMPDock is an adaptation of the RosettaDock (32) protocol that locally searches for docked structures in two stages. First, the algorithm performs a coarse-grained search with random rigid-body rotations and translations. Then, an all-atom stage includes side-chain optimization and minimization along side chain torsion degrees of freedom. RosettaMPDock uses an implicit-membrane approach to evaluate complexes in the context of the lipid bilayer (40, 41) For each of the 19 SERCA-PLB docked complex starting structures we generated 5,000 high-resolution candidate structures. The resulting structures were ranked by PLB angle of rotation around its long axis, tilt angle relative to the membrane normal, and the ΔΔ*G* of binding to SERCA. A summary of the strongest interactions of PLB with each structural state of SERCA is provided in **Table 2** and the full list of SERCA-PLB interactions is provided in Supplementary **Table S2**.

**Table 2:** Summary of top ranking PLB interactions with different SERCA states identified through protein docking

| Enzymatic State | Binding Site | # | Starting Angle (°) | Axial Angle (°) | Energy (REU) | $z_{\text{site}}$ | $z_{\text{all}}$ |
| --- | --- | --- | --- | --- | --- | --- | --- |
| E1-2Ca (1SU4) | M6/M9 | 0 | 11 | 6 | -53.1 | -4.3 | -6.1 |
|  |  | 1 | 46 | 25 | -56.9 | -5.8 | -2.8 |
|  | M3 | 9 | 280 | 273 | -46.2 | -3.5 | -4.5 |
| E2 (3AR4) | M6/M9 | 1 | 319 | 321 | -49.3 | -2.7 | -2.8 |
|  |  | 17 | 89 | 87 | -47.4 | -2.0 | -2.0 |
|  | M3 | 6 | 335 | 334 | -45.1 | -3.0 | -0.9 |
| E1-Like-PLB (4KYT) | M6/M9 | 2 | 347 | 6 | -53.8 | -4.6 | -4.8 |
|  |  | 5 | 277 | 267 | -54.8 | -5.0 | -5.2 |
|  | M3 | 6 | 278 | 298 | -45.2 | -3.0 | -1.0 |

For the E1-like state of SERCA (4KYT), we observed two SERCA-PLB complexes at the M6/M9 site with strong binding affinity with rotational axial angles of 6° and 267°. Both structures exhibit a strong binding funnel (**Fig. 4a**), meaning that the conformations with the lowest ΔΔ*G* of binding are concentrated at a specific PLB axial rotation angle. Further, both z-scores indicated strong and specific interactions, with *z*_all_ of −4.3 for the 6° PLB orientation and −5.2 for the 267° orientation. These docked complexes are represented in **Fig. 4b** in red and blue, respectively. The first orientation was closest to the SERCA-PLB crystal structure (4KYT) with an all-atom root-mean-squared-deviation (RMSD) between PLB from the crystal structure and PLB from the model of 0.89 Å (**Fig. 4b**). **Fig. 4c** shows a helical wheel representation of PLB, highlighting key residues in the interface with SERCA for the 6° (red) and 267° (blue) orientations, and residues that are common to the interfaces of both orientations (orange). The docking results implicating this face of the PLB helix are in harmony with previous biochemical and cross-linking studies (42, 43) and are consistent with the interpretation of the SERCA-PLB co-crystal X-ray structure (10). Global docking also identified a PLB rotational orientation that interacted with the M3 accessory site, though the apparent binding affinity was weaker than those observed for binding to the M6/M9 canonical site (**Supplementary Fig. S5**). M3-binding yielded interface energy of −45.2 Rosetta energy units (REU) and both *z*-scores indicated weaker binding (**Table 2, Supplementary Table S2**). Overall, the binding affinity for the M6/M9 is only modestly better (21% improvement in REU) than the M3 site, suggesting that M3 is a reasonable alternative interface for PLB.

**Figure 4:**
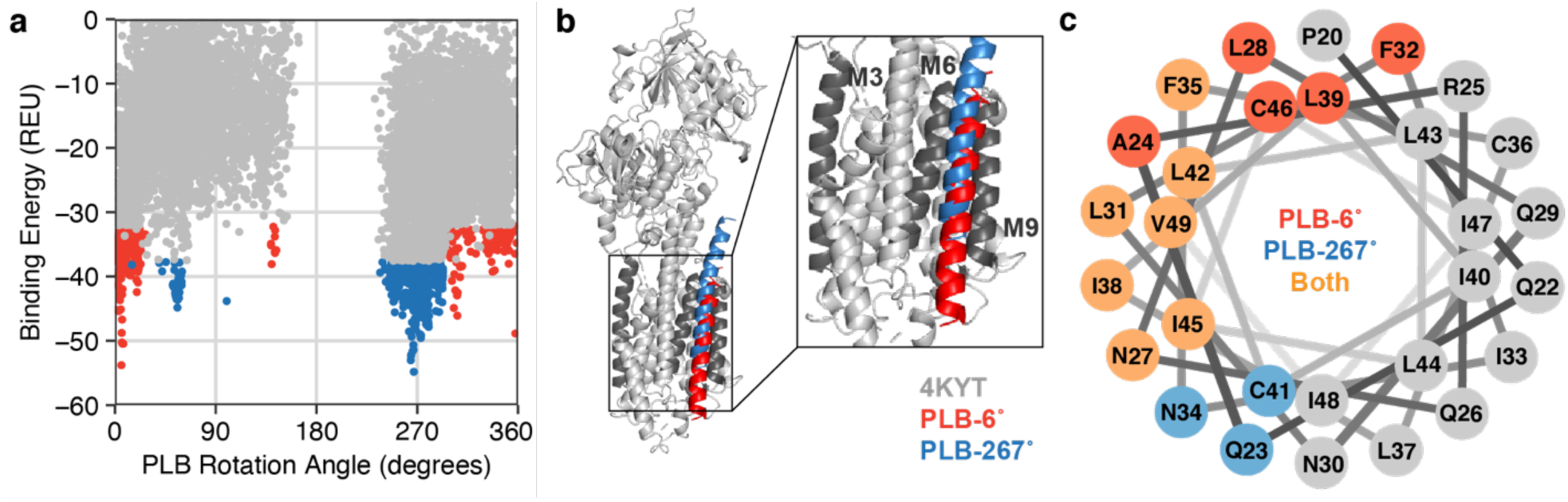
High-resolution model of PLB interaction with the SERCA 4KYT canonical cleft. Molecular docking identified two PLB orientations interacting with the 4KYT canonical cleft, characterized by axial rotation angles of 6° and 267°. (a) Ranking of PLB orientations by axial rotation angle and binding energy. The top 5% scoring points are colored in red for the 6° conformation and blue for the 267° conformation. (b) Conformation of the SERCA-PLB regulatory complex, with a 2x zoomed view of the transmembrane domains. The 6° PLB orientation is shown in red and then 267° orientation is shown in blue, with the M3, M6, and M9 helices highlighted in dark gray. (c) Helical wheel diagram showing the PLB interface residues. Positions unique to the 6° orientation are shown in red, positions unique to the 267° conformation are shown in blue, and mutual positions are colored in orange.

#### Docking of PLB to different enzymatic states of SERCA

The global docking results were also consistent with a multiplicity of interactions between PLB and other conformations of SERCA. Interestingly, there were two strong binding interactions between PLB and the E2 conformation. The first was PLB binding to M6/M9 at 321° with a binding affinity of −49.3 REU (**Supplementary Fig. S6a-c**) and the second was PLB interacting with the alternative M3 site at 334° with a binding affinity of −45.1 REU (**Table 2, Fig. 5a and Fig. 5d**). While the gap between binding affinities to the canonical cleft and M3 site was 10 REU for the E1-like-PLB state (4KYT), the gap narrowed in the E2 (3AR4) state to only 4 REU. Further, the interaction z-scores were suggestive of a significant interaction with this site.

**Figure 5:**
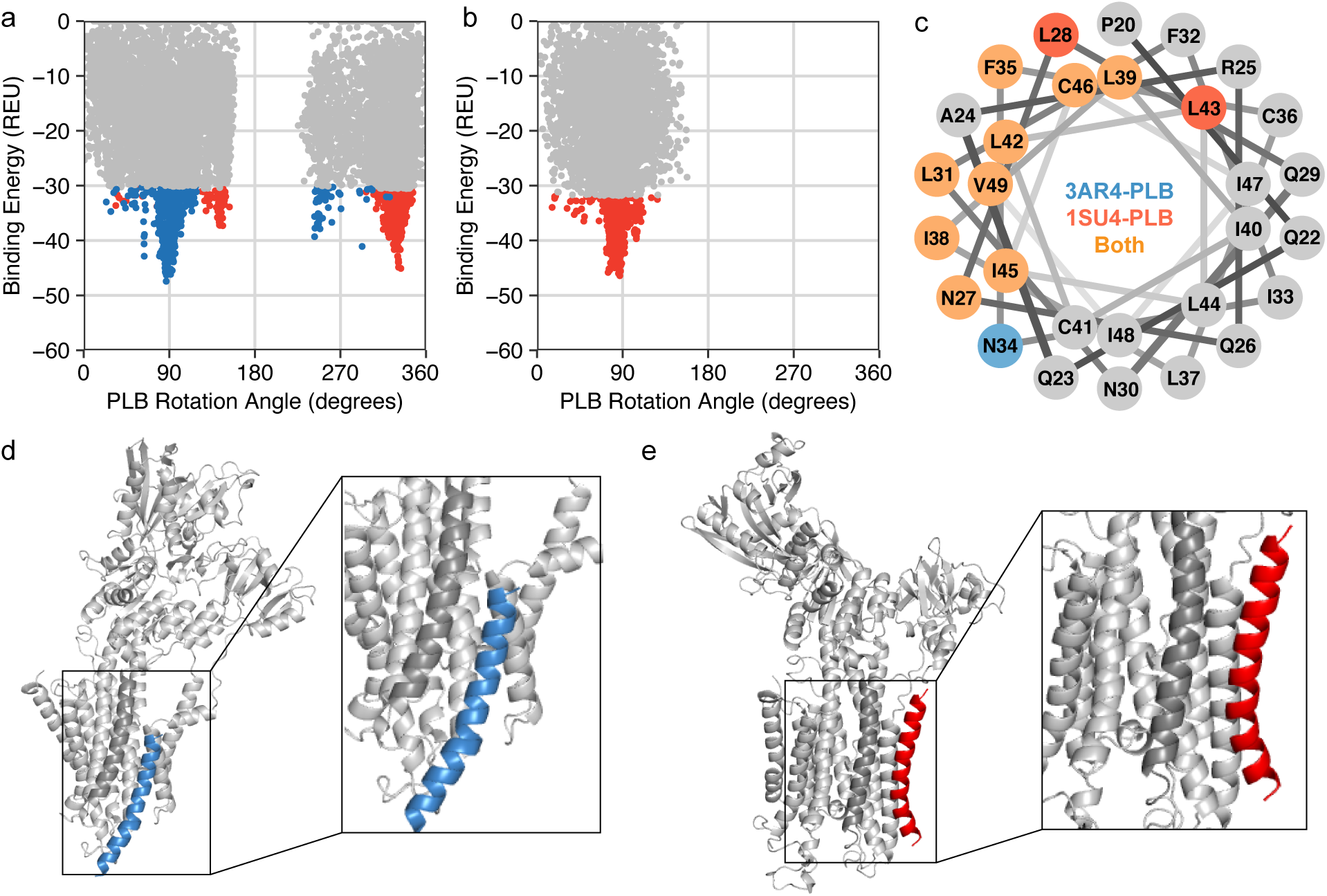
High-resolution models of PLB interaction with the M3 accessory site in the E1 and E2 enzymatic states of SERCA. Molecular docking identified PLB interaction with the M3 helix of the 3AR4 (E2) and 1SU4 (E1-2Ca) enzymatic states of SERCA. Panels (a) and (b) show a ranking of PLB orientations by axial rotation and binding energy, with the top 5% scoring points shown in blue and red for 3AR4 and 1SU4 respectively. (c) Helical wheel diagram showing PLB interface residues: side chains only interacting with 3AR4 are shown in blue, positions only interacting with 1SU4 are shown in red, and mutual positions are shown in orange. Panels (d) and (e) shown structural models for PLB interactions with 3AR4 and 1SU4 respectively. PLB is highlighted in red or blue, and a 2x zoomed representation of the transmembrane domain is shown to the right of the full SERCA model.

Finally, we also examined the affinity of PLB for different sites of the Ca-bound structure of SERCA (1SU4). Overall, we found that PLB bound the most tightly to the M6/M9 canonical site for two rotation angles, 6° and 25° (**Supplementary Fig. S6d-f**). Both sites exhibited strong binding funnels with z_all_ scores of −6.1 and −2.8 respectively. Interestingly, the first conformation was the same binding angle as the low energy E1-like M6/M9 structure, suggesting that face of the helix encodes an important sequence for SERCA interaction. PLB also bound to the M3 site at an angle of 280° (**Fig. 5b,e**), a similar angle for which moderate binding was predicted for docking to the E1-like structure. These data emphasize the role of M3 as a important site for PLB binding. Binding to M3 of SERCA has been observed for both PLB (29) and SLN (44) and is hypothesized to play a role in regulating the maximal activity (V_max_) of SERCA (45, 46).

#### Comparison of WT-PLB and PLB4 Binding to SERCA

To investigate the contribution of specific key sidechains to the energetics of the PLB-SERCA regulatory complex, we repeated the protein-protein docking experiments with the quadruple mutant PLB4 (N27A, N30C, L37A, V49G) that was used to obtain the 4KYT X-ray crystal structure (**Supplementary Table S3**). This superinhibitory mutant binds more avidly than the WT sequence (47). Interestingly, PLB4 docking solutions did not include significant binding to the alternative site on SERCA helix M3. At the canonical M6/M9 site, we observed similar docking for WT-PLB and PLB4, with a narrow binding funnel and interface energies ranging between −54 and −57 REU (**Table S3, Fig. 6a**), but the interface energy was slightly more favorable for docking of PLB4. In addition, there were fewer favorable orientations for PLB4 (**Fig. 6a**) than for WT (**Fig. 4a**). The data suggest that the mutations that enhance PLB binding to SERCA increase the specificity of the interface with regard to the range of favorable axial rotational angles of PLB and the preference for the M6/M9 cleft of SERCA. We noted that the SERCA-binding interface was different from WT, and surprisingly, several of the mutated residues faced the bilayer, rather than the SERCA-PLB interface. Only the N27A and V49G mutant residues participated in direct interactions with SERCA (**Fig. 6b-c**).

**Figure 6:**
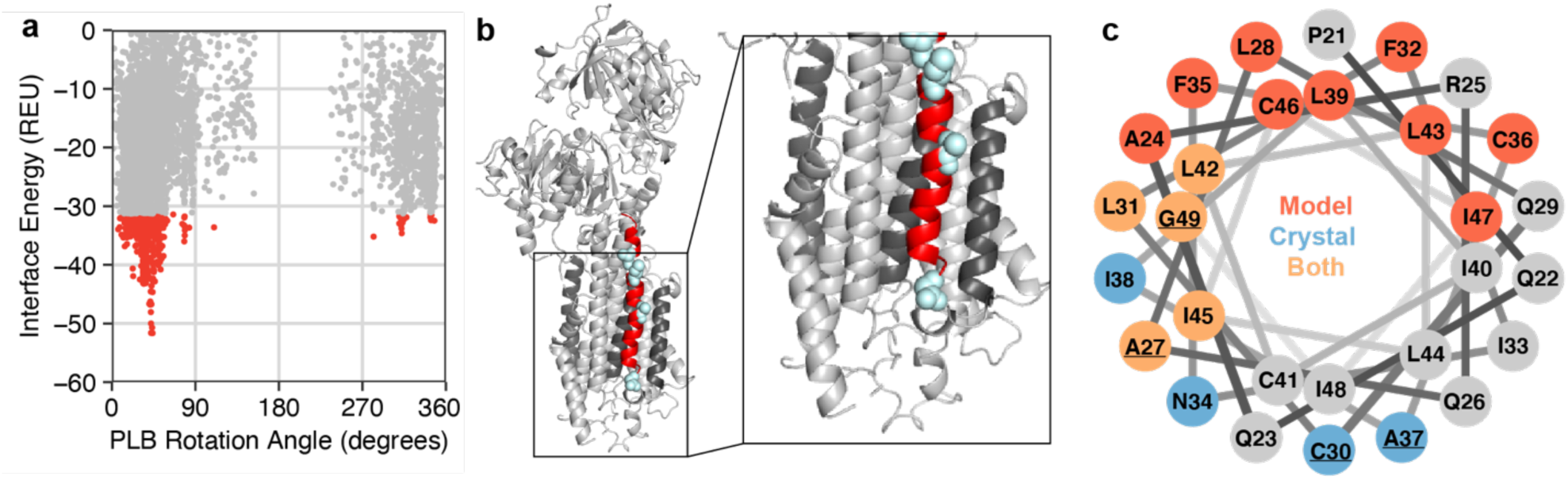
Interactions between SERCA and the PLB4 variant. Protein-protein docking identified interactions between the PLB4 variant (N27A, N30C, L37A, V49G) and the canonical cleft of the E1-Like state of SERCA. Panel (a) shows a ranking of PLB4 orientations by axial rotation and binding energy with the top 5% scoring points shown in red. (b) Structural model of PLB4 interaction with the canonical cleft of 4KYT. PLB4 is highlighted in red with mutation positions colored in light blue. A 2x zoomed representation of the transmembrane domain is shown to the right of the full SERCA model. (c) Helical wheel diagram showing PLB4 interfaces with different enzymatic states of SERCA. The PLB4 mutant sites are underlined. Positions colored in red are part of the docking model interface, positions in blue are part of the crystal structure interface, and positions in orange are common to both.

## Discussion

The goal of this study was to test the hypothesis that PLB may interact with alternative binding sites on SERCA and determine how the population of the alternative sites by PLB may shift with the transporter’s changing structural poise. The motivation for the present computational study comes from consideration of previous physical measurements: a) SERCA-PLB binding affinity changes with SERCA conformational changes (16), but PLB remains bound to SERCA throughout the enzymatic cycle (16, 19, 20); b) cryo-EM studies showed densities attributable to PLB near SERCA TM helix M3 (45, 48); c) X-ray crystallography of an analogous transporter, NKA, showed its cognate regulatory peptide bound to the outside of TM helix M9. We investigated modes of SERCA-PLB interaction through a combined strategy of hypothesis-driven steered molecular dynamics and unbiased protein-protein docking. Overall, the data support the hypothesis for multiple, loosely defined binding sites. The results provide new insight into the diverse modes of interaction for the PLB-SERCA regulatory complex.

### Structural Determinants of the SERCA-PLB Regulatory Complex

The most important point of comparison for the present results is with the high-resolution structure of SERCA co-crystalized with PLB (4KYT) (10). While that structure did not resolve the PLB cytoplasmic domain, it did reveal details of an interface between the PLB transmembrane domain and SERCA. PLB was found at the expected location, in the canonical binding cleft comprising SERCA helices 2, 6, and 9. Residues that were identified as important for the PLB-SERCA interaction in that structure and previous biochemical studies (42, 43) also emerged as key elements in the SMD and unbiased protein-protein docking analysis performed here. Key binding residues identified by biochemistry and crystallography include N27*, N30*, L31, N34, L37*, I38, L42, I45, and V49* (residues marked with asterisks indicate residues mutated to create the superinhibitory mutant PLB4) (10). Those residues were the majority of the interface observed by docking SERCA E1-like conformers with the most favorable PLB rotational orientations (**Fig. 4c**). Those residues also partially overlapped with other PLB binding interfaces observed from protein-protein docking **(Fig. 5c)** and SMD. The other major surface of PLB that repeatedly appeared in docking experiments was the upper left quadrant of the helical wheel **(Figs. 4c, 5c, and 6c)**. Engagement of this surface was frequently associated with a significantly different binding position for PLB, suggesting movement of the regulatory peptide either through an upward shift in the membrane **(Fig. 5d and 5e)**, or translocation to a completely different alternative site (**Fig. 2e**). The involvement of residues near the top of the helical wheel was also favored for the E1 Ca-bound conformation of SERCA (1SU4, **Fig. 5e**), consistent with the concept that SERCA conformational changes alter the nature of the binding of PLB, or, alternatively, that different modes of PLB binding are important for different enzymatic states in the SERCA transport cycle. The data may reconcile apparently contradictory studies of the calcium dependence of chemical cross-linking and FRET experiments. PLB-SERCA FRET is maintained in high Ca, suggesting the complex is intact (16, 19, 20), but cross-linking of PLB is lost in high Ca (11, 12, 43, 49). If the rotational angle and vertical register of PLB changes with the E2-E1 transition it is not surprising that crosslinking of specific residues is greatly diminished.

In this regard, it is instructive to compare the results of docking of wild-type PLB to SERCA with docking of the high affinity, super-inhibitory quadruple mutant PLB4. The latter interacts very avidly with SERCA (11) and provided a sufficiently stable regulatory complex to achieve co-crystallization for X-ray studies (10). Plots of the interface energies of PLB rotational orientations show similar landscapes, with PLB4 funnels (**Fig. 6a**) appearing at similar axial angles as those manifested by WT-PLB (**Fig. 4a**). However, the WT sequence yields more deep funnels (**Fig. 4a** and **Supplementary Fig. S4**), suggesting more energetically acceptable docking options. Therefore, we hypothesize that PLB mutations that caused tighter binding to SERCA (42) and yielded well-ordered structures for crystallization (10) tailored PLB to bind in a specific orientation to a specific (canonical) binding site of a specific SERCA conformer.

### Alternative Modes of Binding of PLB to SERCA

Protein docking studies identified transmembrane segment M3 in both the E1-2CA and E2 states of SERCA as a potential interaction interface. Of the docked complexes between SERCA and the wild-type PLB monomer, one of the solutions (model #6) is similar to the previously described MD simulations of the SERCA-PLB pentamer complex based on electron crystallography (29). The docked complex is not identical to the previous complex, though this is likely due to the PLB pentamer used in the prior docking and MD simulations. In addition, that previous study used the human PLB sequence (in which residue 27 is a lysine), while the present docking analysis follows the X-ray crystallography study in using the canine PLB (with an asparagine at position 27). Nonetheless, the PLB monomer in the docked complex (**Fig. 5**) interacts with M3, but it is located more toward transmembrane segment M1 of SERCA and it is shifted upward by one turn of the helix. The interaction interface between SERCA and PLB is similar in the two models, with key residues such as F32 and I45 of PLB and K26 and Y272 of SERCA contributing. In the docked complex, N34 interacts with E258. This fits with the previous conclusion that electrostatic interactions, at least in part, draw PLB into this region of SERCA. The negatively charged residues include D254 and E255 on M3 and additional residues on M1 (e.g. D59 and E58). Finally, it is interesting to note that the protein docking of the PLB4 variant used in the X-ray crystal structure of the SERCA-PLB complex (10) did not yield a docked complex at the M3 accessory site of SERCA. Only the wild-type PLB yielded satisfactory docking to M3 of SERCA.

### Complementary Approaches for Interrogating Membrane Protein Complexes

Steered molecular dynamics and protein-protein docking provided complementary perspectives in evaluating PLB-SERCA binding interactions. SMD offers an all-atom view of SERCA’s behavior in a biologically realistic membrane, quantifying rupture forces as an index of protein-protein binding affinity. On the other hand, protein-protein docking provides an unbiased strategy. RosettaMPDock, in particular, is one of a few specialized docking methods that can efficiently and inexpensively explore new conformations while considering the physical properties of the surrounding lipid bilayer. Overall, the results from the two types of experiments were in harmony. Most notably, both suggested that the canonical binding cleft is the most favorable site for PLB binding.

On the other hand, the methods provided different insights into the interactions between PLB and alternative sites, and the binding of PLB to different SERCA conformers. Both SMD and protein-protein docking simulations suggest that M3 is a plausible alternative site. Docking yielded a favorable PLB orientation at 298° binding with weak-to-moderate affinity (**Fig. 5**). By SMD, the best M3 binding is seen for the 180° and 90° orientations, yielding a rupture force that was 1 or 2 standard deviations from the mean rupture force, respectively. Interestingly, the 180° orientation that showed relatively weak affinity by SMD (1 standard deviation above the mean), and did not appear among high-scoring docking results, suggesting it is not a preferred orientation. In contrast, there was an 80° PLB orientation with weak binding affinity (−35 REU). We regard the 80° value (from docking) and 90° value (from SMD) as representing a similar possible orientation of PLB bound to M3. Another difference between the experiments is that the 298° orientation observed in docking was not tested by SMD, because those experiments systematically evaluated fixed PLB orientations at 90° intervals. The identification of a favorable “in-between” orientation at 298° demonstrates the added value of an unbiased docking strategy for identifying new quaternary arrangements that may be important for regulating SERCA transport function.

Conversely, SMD provided insight into a possible interaction that was not detected by docking. SMD experiments showed binding to hypothetical site on the outside of M9 was relatively strongly site- and orientation-specific for PLB in for the E1-like Ca-free (3AR4) conformation of SERCA. The 180° and 270° orientations of PLB bound to M9 yielded rupture forces that appeared equally favorable to the canonical cleft. However, despite this apparent site selectivity, it was a moderately weak binder overall, as compared to all other interactions. Thus, the site was not ranked highly enough to be discovered by protein-protein docking. On the basis of the SMD experiments, we suspect that the outside of M9 may be populated by PLB for some conformations of SERCA, such as the E2 Ca-free state represented by 3AR4 (**Fig. 2**). An analogous interaction is that of the sodium/potassium ATPase, which binds FXYD proteins on the outside of M9 of the NKA alpha (catalytic) subunit. This regulatory complex also benefits from the contribution of additional contacts between the FXYD extracellular residues with the NKA beta subunit (26), accounting for the apparently stable occupation of that position in the crystal structure. Overall, the disparity between the SMD and docking results underscores the value of complementary approaches for evaluating metastable interactions that may be biologically significant.

### Challenges for modeling of protein-protein interactions in the membrane

In this study, we applied two complementary strategies to investigate a physiologically important integral membrane interaction. This dual strategy enabled the computational feasibility of exploring the wide protein-protein interaction space for a large target (>1,000 residues) while also considering the context of a heterogeneous membrane environment. This is an exciting study in membrane protein structural biology because specialized methods for exploring membrane protein-protein interactions are still in their infancy. In particular, there are many open questions about the biophysical questions driving protein-protein interactions in the nonpolar membrane environment. With this, these specialized methods require improvement in several areas. For instance, the protein-protein docking program kept the backbone of both SERCA and PLB fixed. However, it is likely that PLB bends or straightens to enhance shape complementarity with its binding partner. This is especially notable because transmembrane membrane helices are notorious for kinks and curvature (50). Programs for incorporating backbone flexibility have emerged for soluble proteins (51, 52) and these developments will soon translate to membrane proteins. Another challenge is incorporating the effect of bilayer deformations induced by the protein that influence the landscape of available protein-protein complex conformations (53). Further, the all-atom membrane in SMD only included one lipid type, whereas biological membranes include hundreds of different lipid types (54). While building membrane models complex lipid compositions is computationally expensive due to a long equilibration time, improved computing capabilities will make these models within reach.

### Summary

The conclusion that emerges from the present results is that the PLB-SERCA binding interactions are more diverse than can be appreciated from previous biochemical and structural biology studies that suggested PLB binds with a fixed orientation to a single site on a specific SERCA conformer. Instead, the present data are consistent with spectroscopic results that suggest PLB interacts with all SERCA enzymatic states and structural data that suggest additional alternative binding sites. We propose PLB equilibrates between different modes of binding. Some of these mode changes are minor shifts of PLB within a binding pocket (e.g. rotation about the PLB long axis, or normal translations in the bilayer), while others are translocation to completely different sites (e.g. to the outside of M9 or to M3). Based on the increased apparent specificity of the superinhibitory PLB4 mutant for the M6/M9 site, we conclude that the canonical binding mode mediates functional inhibition, and we presume that the other sites are non-inhibitory, or possibly even stimulatory (45, 46, 55). Thus, PLB transitions between different structural poises may deliver different functional outcomes as is appropriate for the varying demands of different physiological conditions.

## Methods

### Steered molecular dynamics simulations

The SERCA/PLB complex was modeled from crystal structures of SERCA1a, the skeletal muscle isoform for which many conformations are available (24). This isoform has high homology to the cardiac specific Ca pump, SERCA2a, and the recently published first structure of SERCA2a showed that the known PLB-binding site is highly conserved between these isoforms (56). Here we generated models starting from three different SERCA1a structures: 1SU4 (36), representing an E1 Ca-bound state; 3AR4 (35), representing an E2 Ca-free state; and 4KYT (10), representing a Ca-free state bound to PLB. The latter Ca-free X-ray crystal structure was designated an E2 structure, but the conformation resembles an E1 structure, and is referred to as “E1-like” in the present study. In that x-ray crystal structural solution, the cytoplasmic domain of PLB was not observed. In addition, 4KYT contains structure of mutated PLB; for experiments that simulate interaction of WT-PLB with SERCA we mutated the residues back to the native sequence. In SMD experiments, we examined transmembrane domain residues 22-52 docked in the canonical binding cleft of 4KYT or the equivalent site on 1SU4 or 4AR4. To test the possibility of PLB binding to the putative alternative sites on SERCA, we created a model of the PLB bound to M9 using as a guide the NKA co-crystal structure with a FXYD protein(26, 27). PLB was also docked to M3 to test a possible interaction hypothesized from cryo-EM studies of SERCA-PLB co-crystals (30, 45).

### Protein-protein docking simulations

A search for SERCA-PLB interaction sites was performed using global and local docking. Each crystal structure was first refined using RosettaMPRelax (39) to erase artifacts from crystallization and prior binding ligands. To accomplish this, the protocol performs cycles of small backbone torsion moves followed by side-chain repacking and energy minimization of all torsion angles (*ϕ, ψ, χ*)(57). For each starting structure, 50 PLB orientations were generated, and the lowest energy conformation was used as the next starting pose. Next, the ClusPro Fast-Fourier Transform (FFT)-based rigid-body docking server (38, 58) was used to perform a global search for SERCA-PLB binding sites. Then, the following criteria were used to filter solutions from the best-scoring clusters: (1) the conformers are in the correct direction (e.g. V49G) and (2) the conformers span the membrane.

Following global docking, RosettaMPDock (39) was used to locally search for PLB binding orientations. RosettaMPDock an adaptation of the RosettaDock (32) minimization protocol that locally searches for docked PLB orientations through two stages (a) a coarse-grained stage to quickly identify favorable orientations and (b) an all-atom refinement stage that optimizes the rigid body position and side-chain rotamers. To account for the membrane environment, RosettaMPDock additionally samples the protein-membrane orientation and uses an implicit-membrane energy function to evaluate complexes in the context of the lipid bilayer (40, 41). For each starting pose, 5,000 PLB orientations were generated. Then, the interface analyzer protocol.(59) was used to compute two properties: (a) *Axial Angle* – the angle between the second principal axis of the 4KYT PLB conformation and the candidate PLB conformation and (b) ΔΔ*G*_bind_ – the change in Rosetta energy when the chains are separated versus when they are in complex.

### Z-score analysis

To quantify the significance of each major mode of SERCA-PLB binding, we computed two scores that measure the similarity of binding conformations, with values of zero indicating the conformations are identical, and larger values corresponding to greater differences. The first score called *z*_all_ evaluates the relationship of a particular binding conformation relative to the mean of all possible binding conformations. The second score, called *z*_site_, computes the same quantity but relative to the mean of all structures interacting with a specific site (e.g. M3, M6/M9 or the outside of M9). For docking, the mean was computed from the top 5% of structures ranked by ΔΔ*G* of binding.

## Acknowledgements

This work was supported by the Loyola University Chicago Cardiovascular Research Institute. R.F.A. was funded by a Hertz Foundation Fellowship and an NSF Graduate Research Fellowship. J.J.G. was supported by NIH GM-078221. S.L.R and H.S.Y. were supported by NIH HL-092321 and HL-143816.

## Supplementary Material

*Supplementary figures, tables and methods can be found in SupportingInformation*.*pdf*.

