## Supporting Information for "Protein Docking and Steered Molecular Dynamics Reveal Alternative Regulatory Sites on the SERCA Calcium Transporter"

Rebecca F. Alford[1], Nikolai Smolin[2], Howard S. Young[3], Jeffrey J. Gray[1], and Seth L. Robia[2#]
[1] Dept. of Chemical & Biomolecular Engineering, Johns Hopkins University, Baltimore MD
[2] Dept. of Cell and Molecular Physiology, Stritch School of Medicine, Cardiovascular Research Institute, Loyola University Chicago, Maywood, IL
[3]Dept. of Biochemistry, University of Alberta, Edmonton, Alberta, Canada

### Supplementary Methods

#### *Model preparation*

The structure of each enzymatic state of SERCA was downloaded from the Orientations of Proteins in Membranes Database (OPM: https://opm.phar.umich.edu/). Protein structures from this database are transformed into a reference membrane coordinate frame and the orientation is optimized using an anisotropic membrane model. PDB files were cleaned and renumbered using the `clean_pdb.py` script in `Rosetta/tools/protein_tools/scripts`. The following PDB codes were used: PDB model 4KYT for the E1-like-PLB state, 3AR4 for the E2 state, and 1SU4 for the E1-2Ca state. The transmembrane spanning topology was defined from structure using the `span_from_pdb` application.

#### *All-atom refinement of SERCA starting structures*

All-atom refinement of all three SERCA states was performed to remove steric clashes, crystallization artifacts, and binding history (in the case of 4KYT). Refinement was performed with the RosettaMPRelax application using the command line below.

```
Rosetta/main/source/bin/rosetta_scripts.linuxgccrelease
      -parser:protocol membrane_relax.xml      # Path to Rosetta script
      -in:file:s 4KYT.pdb                      # Input PDB file
      -nstruct 50                              # Generate 10 models
      -mp:setup:spanfiles 4KYT.span            # Input spanfile
      -mp:scoring:hbond                        # Hbond potential
      -packing:pack_missing_sidechains 0       # Wait to pack sidechains
                                               # until membrane is present
      -parser:script_vars
         -sfxn_weights=mpframework_smooth_fa_2012# Weights
      2012
```

Here, `membrane_relax.xml` is a RosettaScript defining an all-atom refinement protocol in the context of the membrane. The contents of `membrane_relax.xml` are listed below.

```
<ROSETTASCRIPTS>
      <SCOREFXNS>
            <ScoreFunction name="memb_hires" weights="%%sfxn_weights%%" />
      </SCOREFXNS>
      <MOVERS>
            <AddMembraneMover name="add_memb"/>
            <MembranePositionFromTopologyMover name="init_pos"/>
            <FastRelax name="fast_relax" scorefxn="memb_hires" repeats="8"/>
      </MOVERS>
      <PROTOCOLS>
            <Add mover="add_memb"/>
            <Add mover="init_pos"/>
            <Add mover="fast_relax"/>
      </PROTOCOLS>
      <OUTPUT scorefxn="memb_hires" />
</ROSETTASCRIPTS>
```

#### *ClusPro Global Docking calculations*

To generate global docking models, we used the ClusPro server (https://cluspro.bu.edu/). The SERCA chain was designated as the receptor and the PLB chain was designated as the ligand. No additional filters or masks were applied during docking.

#### *Rosetta membrane protein-protein docking simulations*

To refine each global docking model in context of the membrane, we used the RosettaMPDock local docking algorithm. First, we ran the prepack protocol with the following application:

```
Rosetta/main/source/bin/docking_prepack_protocol.linuxgccrelease
      -in:file:s 4KYT_AB.pdb              # Input PDB with both partners
      -nstruct 1                          # Generate one model
      -score:weights mpframework_docking_fa_2015.wts # Use mp score function
      -mp:setup:spanfiles 4KYT_AB.span    # Input spanfile
      -mp:scoring:hbond                   # Turn on membrane hydrogen bonding
      -packing:pack_missing_sidechains 0  # Wait to pack sidechains until
                                          # membrane is present
```

The MEM residue was removed from the output model. Then, the output model was used as input to the protein-protein docking protocol. The following application and options were used:

```
Rosetta/main/source/bin/mpdocking.linuxgccrelease
      -in:file:s 4KYT_AB_ppk.pdb          # Pre-packed input structure
      -nstruct 5000                       # Generate 1000 models
      -score:weights mpframework_docking_fa_2015.wts # Score function
      -mp:setup:spanfiles 4KYT_AB.span    # Input spanfile
      -mp:scoring:hbond                   # Turn on membrane hydrogen bonding
      -docking:partners A_B               # Partners to dock
      -docking:dock_pert 3 8              # Magnitude of perturbation
      -packing:pack_missing_sidechains 0  # Wait to pack sidechains until
                                          # membrane is present
```

5000 models were generated with a default docking perturbation of 3 Å translation and 8 degrees rotation. Next, the models were rescored using the Interface Analyzer application with the command line given below.

```
Rosetta/main/source/bin/InterfaceAnalyzer.linuxgccrelease
      -in:file:l 4KYT_AB_models.list      # List of docked models
      -out:file:score_only                # Do not generate PDB files
      -interface A_B                      # Docking partners
```

Finally, we computed the axial rotation angle of PLB using the `calc_micropeptide_angle` application with the command line below.

```
Rosetta/main/source/bin/calc_micropeptide_angle.linuxgccrelease
      -in:file:l 4KYT_AB_models.list      # List of docked models
      -out:file:score_only                # Do not generate PDB files
      -reference_pose PLB_angle0.pdb      # Structure of PLB from 4KYT
```

### Supplementary Tables

**Table S1:** Summary of *z*-scores for PLB conformations explored by steered molecular dynamics

| Enzymatic State | Binding Site | Angle (˚) | Pulling Force (kJ/mol/nm) | $z_{site}$ | $z_{all}$ |
|---|---|---|---|---|---|
| E1-Like-PLB (4KYT) | M3 | 0 | 546 | -1.13 | -0.68 |
| | | 90 | 526 | -1.26 | -1.01 |
| | | 180 | 485 | -1.52 | -1.68 |
| | | 270 | 550 | -1.1 | -0.61 |
| | M6/M9 | 0 | 1292 | 3.73 | 2.77 |
| | | 90 | 982 | 1.71 | 0.76 |
| | | 180 | 838 | 0.77 | -0.18 |
| | | 270 | 914 | 1.27 | 0.32 |
| | M9 | 0 | 686 | -0.21 | -0.25 |
| | | 90 | 731 | 0.08 | 0.36 |
| | | 180 | 688 | -0.2 | -0.22 |
| | | 270 | 616 | -0.67 | -1.2 |
| E2 (3AR4) | M3 | 0 | 544 | -1.14 | -0.71 |
| | | 90 | 640 | -0.51 | 0.86 |
| | | 180 | 698 | -0.14 | 1.8 |
| | | 270 | 572 | -0.96 | -0.25 |
| | M6/M9 | 0 | 848 | 0.84 | -0.11 |
| | | 90 | 814 | 0.62 | -0.33 |
| | | 180 | 729 | 0.07 | -0.89 |
| | | 270 | 720 | 0.01 | -0.95 |
| | M9 | 0 | 658 | -0.4 | -0.63 |
| | | 90 | 580 | -0.9 | -1.7 |
| | | 180 | 811 | 0.6 | 1.47 |
| | | 270 | 838 | 0.77 | 1.84 |
| E1-2Ca (1SU4) | M3 | 0 | 606 | -0.74 | 0.3 |
| | | 90 | 655 | -0.42 | 1.1 |
| | | 180 | 629 | -0.59 | 0.68 |
| | | 270 | 601 | -0.77 | 0.22 |
| | M6/M9 | 0 | 871 | 0.99 | 0.04 |
| | | 90 | 775 | 0.36 | -0.59 |
| | | 180 | 833 | 0.74 | -0.21 |
| | | 270 | 766 | 0.31 | -0.65 |
| | M9 | 0 | 682 | -0.24 | -0.3 |
| | | 90 | 681 | -0.25 | -0.31 |
| | | 180 | 736 | 0.11 | 0.44 |
| | | 270 | 749 | 0.2 | 0.61 |

**Table S2:** Summary of PLB binding conformers in complex with different states of SERCA

| Enzymatic State | Binding Site | # | Starting Angle (˚) | Axial Angle (˚) | Energy (REU) | $z_{site}$ | $z_{all}$ |
|---|---|---|---|---|---|---|---|
| E1-2Ca (1SU4) | M6/M9 | 0 | 11.41 | 5.85 | -53.091 | -4.26 | -6.13 |
| | | 1 | 45.57 | 25.15 | -56.9 | -5.82 | -2.83 |
| | | 2 | 250.55 | 266.67 | -49.41 | -2.77 | -2.19 |
| | | 3 | 35.53 | 55.75 | -47.93 | -2.15 | 0.58 |
| | | 5 | 298.32 | 303.334 | -41.64 | -0.42 | -1.52 |
| | M3 | 9 | 279.61 | 272.93 | -46.15 | -3.49 | -4.45 |
| E2 (3AR4) | M6/M9 | 1 | 319.01 | 321.477 | -49.33 | -2.72 | -2.8 |
| | | 2 | 336.29 | 341.806 | -44.7 | -0.83 | -0.77 |
| | | 17 | 88.58 | 86.711 | -47.43 | -1.95 | -1.97 |
| | M3 | 6 | 335.41 | 333.569 | -45.09 | -2.99 | -0.94 |
| | | 10 | 142.1 | 131.42 | -41.31 | -1.57 | -0.72 |
| E1-Like-PLB (4KYT) | M6/M9 | 0 | 49.82 | 52.63 | -46.23 | -1.46 | -1.44 |
| | | 1 | 29.93 | 36.52 | -43.77 | -0.45 | -0.36 |
| | | 2 | 346.97 | 5.734 | -53.8 | -4.55 | -4.76 |
| | | 3 | 57.03 | 287.873 | -47.82 | -2.11 | -2.14 |
| | | 4 | 252.61 | 255.553 | -44.2 | -0.63 | -0.54 |
| | | 5 | 276.54 | 266.95 | -54.83 | -4.97 | -5.22 |
| | M3 | 6 | 277.7 | 298.45 | -45.2 | -3.03 | -0.98 |
| | | 7 | 292.29 | 69.44 | -41.45 | -1.61 | 0.66 |

**Table S3:** Summary of PLB4 binding conformers in complex with different states of SERCA

| Enzymatic State | Binding Site | # | Starting Angle (˚) | Axial Angle (˚) | Energy (REU) | $z_{site}$ | $z_{all}$ |
|---|---|---|---|---|---|---|---|
| E1-2Ca (1SU4) | M6/M9 | 1 | 17.47 | 17.501 | -53.77 | -3.15 | -2.97 |
| | M3 | 2 | 102.32 | 56.8 | -42.49 | -0.75 | -0.04 |
| E2 (3AR4) | M6/M9 | 1 | 322.12 | 324.08 | -49.53 | -2.289 | -1.87 |
| | | 2 | 337.69 | 338.96 | -49.46 | -2.28 | -1.85 |
| | | 3 | 57.17 | 59.62 | -52.76 | -3.57 | -2.71 |
| | M3 | 4 | 115.47 | 329.37 | -38.75 | 0.283 | 0.92 |
| E1-Like-PLB (4KYT) | M6/M9 | 1 | 25.25 | 24.49 | -56.62 | -3.73 | -3.71 |
| | | 2 | 28.71 | 43.8 | -51.64 | -2.718 | -2.41 |
| | M3 | 3 | 60.93 | 65.29 | -44.91 | -1.41 | -0.674 |
| | | 4 | 55.25 | 52.68 | -45.2 | -1.49 | -0.75 |

### Supplementary Figures

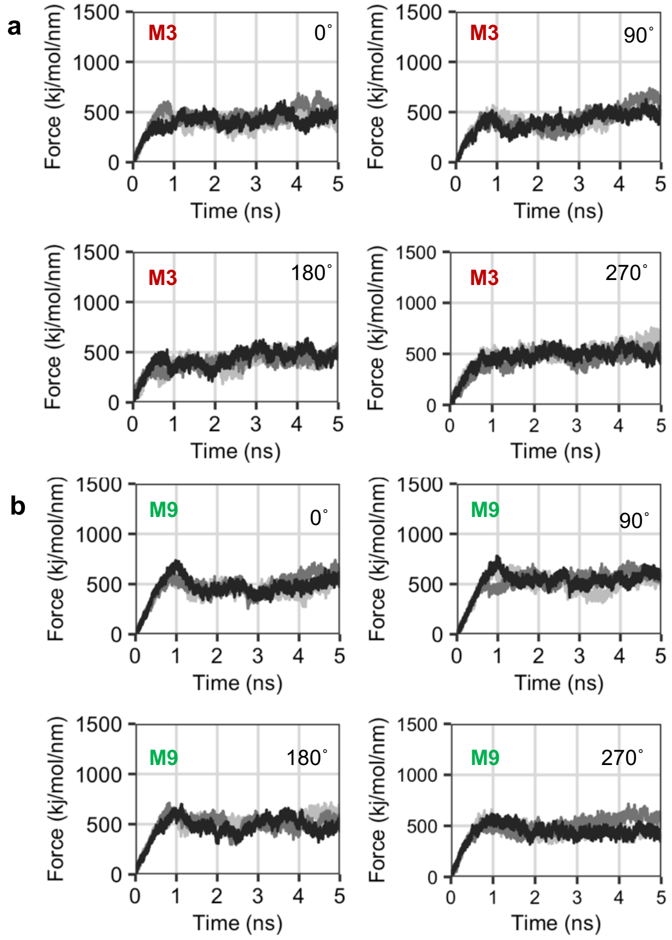

**Figure S1: SMD simulations of the E1-Like-PLB state of SERCA with PLB.** Quantification of force that developed as PLB was pulled from the canonical cleft of the E1-like-PLB state of SERCA (represented by 4KYT). Data are three repeated measures of force for the original orientation of the X-Ray crystal structure (PDB 4KYT) and after axial rotation of PLB by 90˚, 180˚, and 270˚. The data are shown for interaction with the M3 (a) and M9 (b) helices of SERCA.

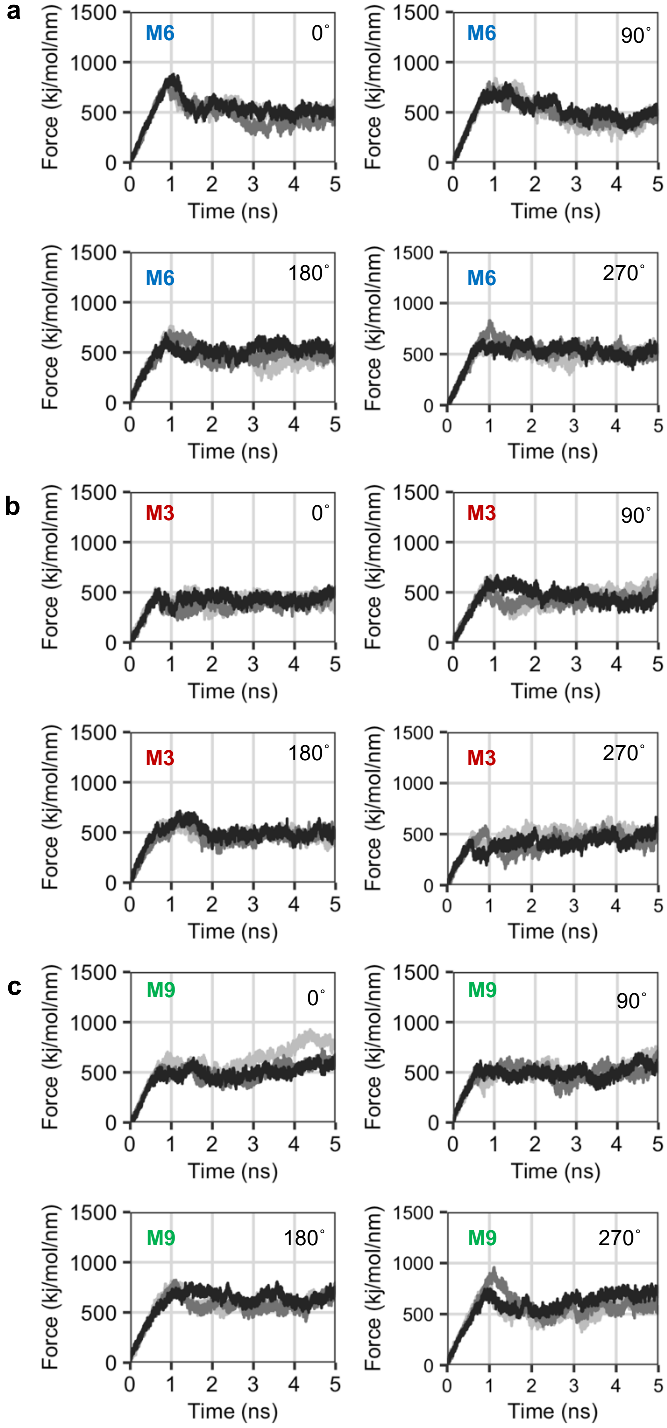

**Figure S2: SMD simulations of the E2 state of SERCA with PLB.** Quantification of force that developed as PLB was pulled from the canonical cleft of the E2 state of SERCA (represented by 3AR4). Data are three repeated measures of force for the original orientation of the X-Ray crystal structure (PDB 4KYT) and after axial rotation of PLB by 90˚, 180˚, and 270˚. The data are shown for interaction with the M6/M9 (a) M3 (b) and M9 (c) helices of SERCA.

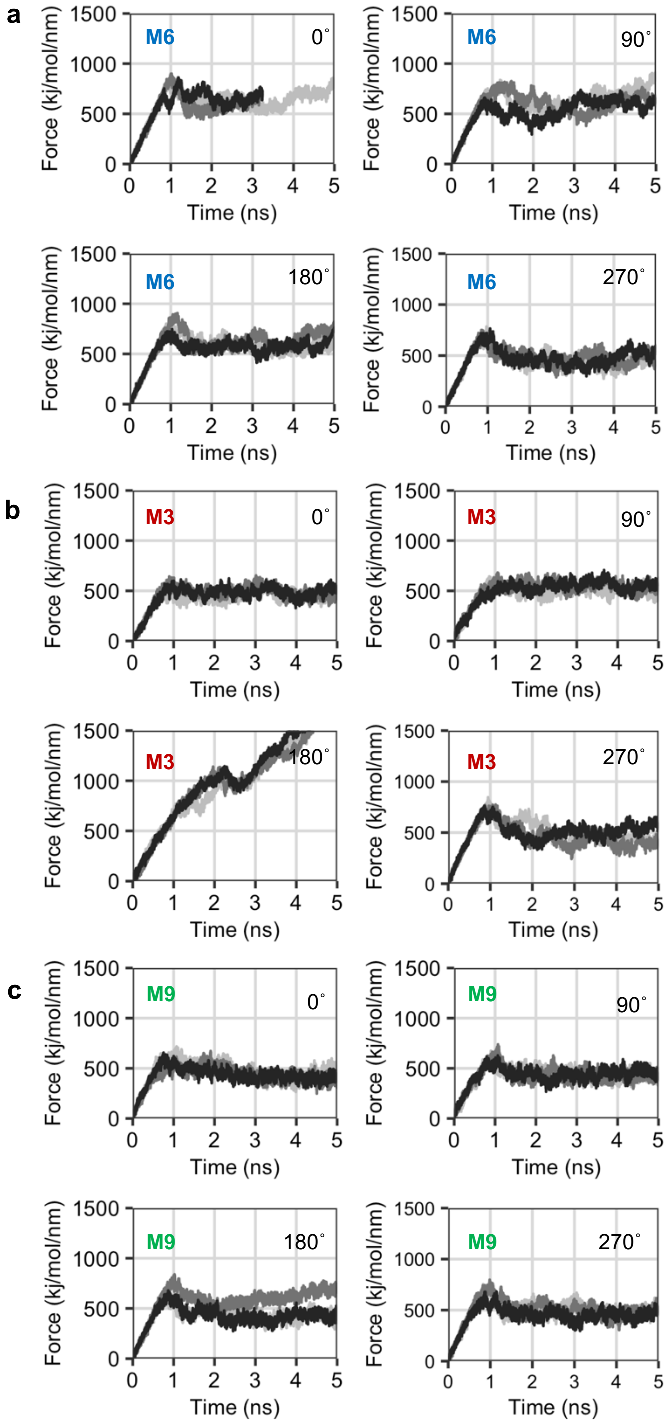

**Figure S3: SMD simulations of the E1-2Ca state of SERCA with PLB.** Quantification of force that developed as PLB was pulled from the canonical cleft of the E1-2Ca state of SERCA. Data are three repeated measures of force for the original orientation of the X-ray crystal structure (PDB 4KYT) and after axial rotation of PLB by 90˚, 180˚, and 270˚. The data are shown for interaction with the M6/M9 (a) M3 (b) and M9 (c) helices of SERCA.

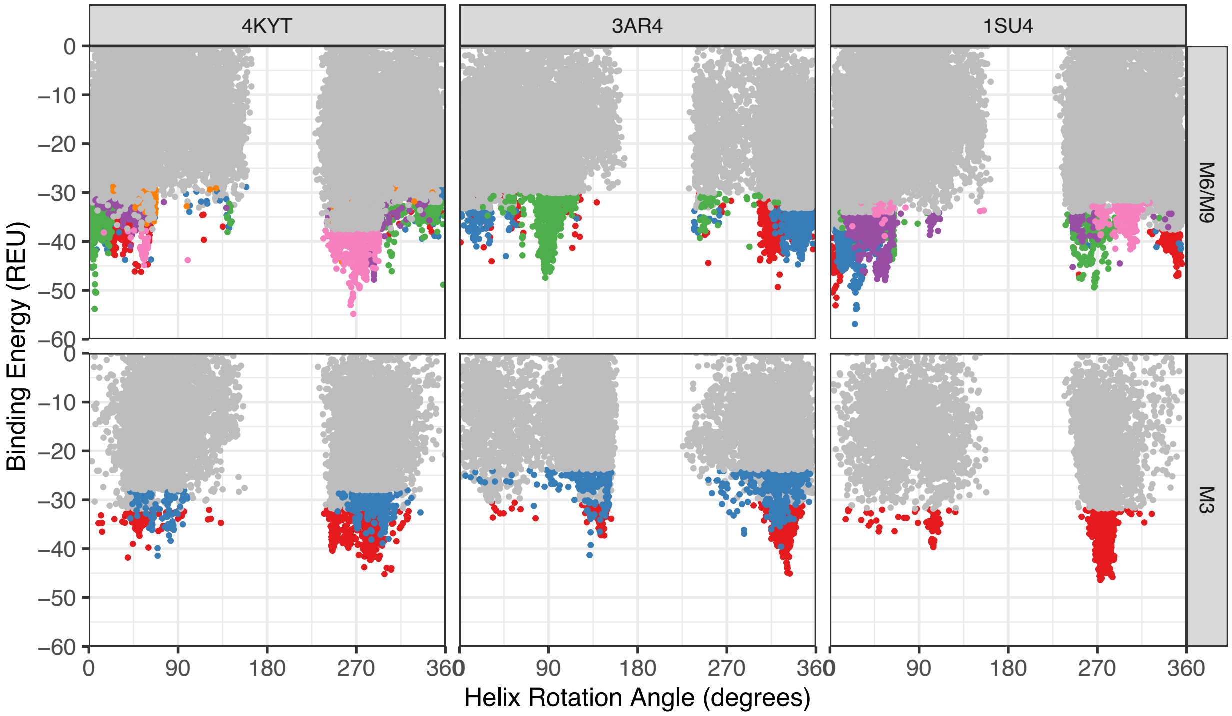

**Figure S4: Ranking of all PLB conformers from all global docking solutions.** Each SERCA-PLB complex model is ranked by axial rotation and binding energy, with each sub-panel corresponding to a particular enzymatic state (4KYT, 3AR4, or 1SU4) and binding site (canonical cleft M6/M9 or M3). The top 5% scoring points are shown in red, blue, orange, green, pink or purple. A different color indicates the model was generated from a different global docking solution.

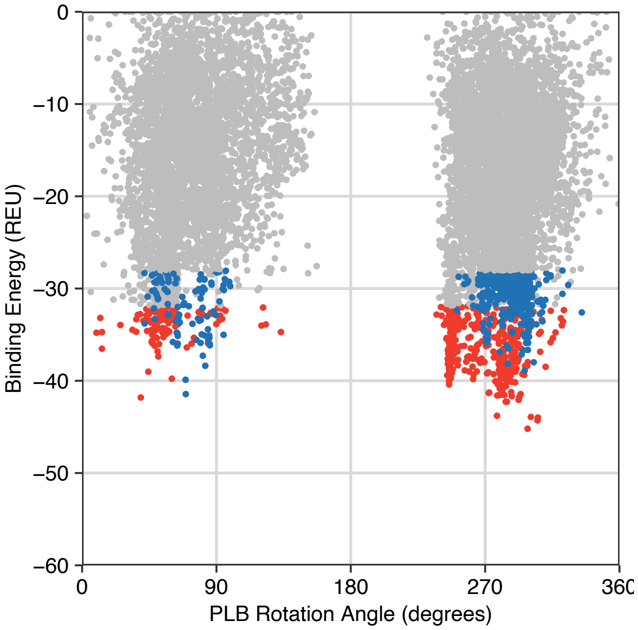

**Figure S5: Ranking of all PLB conformers from docking the E1-Like-PLB state of SERCA with the M3 accessory site.** Each SERCA-PLB complex model is ranked by axial rotation and binding energy and the top 5% scoring points are shown in red or blue. A different color indicates the model was generated from a different global docking solution.

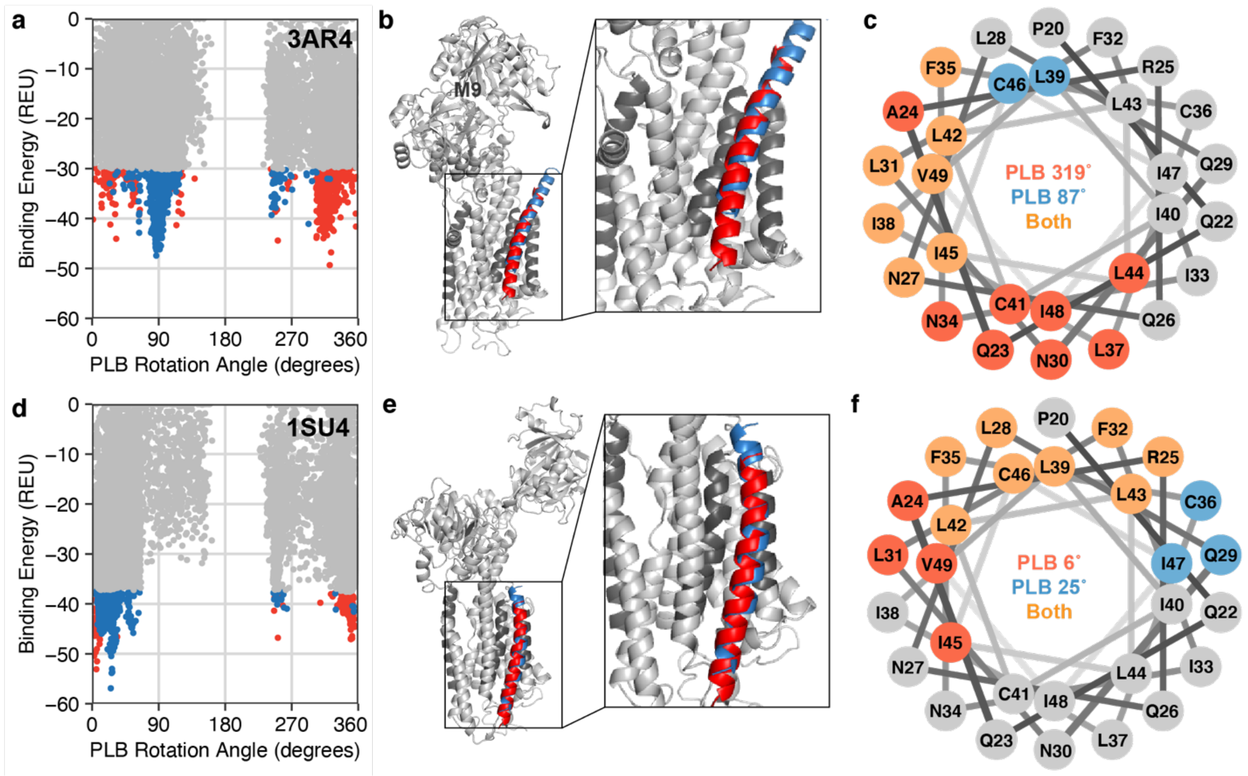

**Figure S6: High resolution models of PLB interactions with the canonical cleft of the E2 and E1-2Ca states of SERCA.** The model of the E2 state (represented by 3AR4) is shown in the top row and the model of the E1-2Ca state (represented by 1SU4) is shown in the bottom row. (a,d) Ranking of each SERCA-PLB complex model by axial rotation and binding energy. The top 5% scoring points are shown in red or blue, with a different color indicating the model was generated from a different global docking solution. (b,e) High resolution model of two possible PLB conformers interacting with SERCA shown in red and blue, corresponding to the colors in panels (a,d). A 2x zoomed view of the transmembrane domain is shown to the right of the SERCA model. (c,f) Helical wheel demonstrating PLB conformer interface residues. Positions colored in red correspond to the first conformer, positions in blue correspond to the second conformer, and mutual positions are in orange.
